# Longitudinal tractography monitors cortical stroke-induced thalamocortical degeneration and neuroprotection by human iPSC-derived neurons

**DOI:** 10.1101/2025.03.17.643730

**Authors:** Sopiko Kartsivadze, Yu-Ping Shen, Linda Jansson, Lucía Galán-Pintado, Keshav Kher, Sara Avilés-Carpintero, Michael Gottschalk, Aref Kalantari, Daniel Mertens, Raquel Martinez-Curiel, Markus Aswendt, Olle Lindvall, Sara Palma-Tortosa, Zaal Kokaia

## Abstract

Cortical ischemic stroke can trigger secondary neurodegeneration in remote brain regions connected to the primary lesion, particularly the thalamus. Although secondary thalamic degeneration is well established, the temporal relationship among early disruption of the thalamocortical pathway, neuronal degeneration, neuroinflammation, and delayed thalamic atrophy remains incompletely defined. Here, we investigated the longitudinal progression of secondary thalamic degeneration after cortical stroke, with particular emphasis on diffusion MRI tractography to monitor changes in anatomically defined thalamocortical pathways. We further determined whether early intracortical transplantation of human induced pluripotent stem cell (hiPSC)-derived neuronal progenitors could protect the remote thalamus and preserve thalamocortical connectivity.

Cortical ischemic stroke was induced by distal middle cerebral artery occlusion in rats, and animals were assessed at multiple time points up to 6 months after stroke using longitudinal volumetric MRI and diffusion MRI tractography, combined with histological analyses of neuronal degeneration and microglial activation. The cortical infarct was established within 3-4 days, whereas alterations in diffusion metrics of thalamocortical pathways were detected within the first days after stroke. Neuronal degeneration in the ipsilateral ventral posterior nucleus (VPN) was evident at 2 weeks and preceded measurable VPN atrophy, which began at 3 months. Microglial activation also peaked at 2 weeks, coinciding with the onset of neuronal degeneration. Early intracortical transplantation of cortically primed hiPSC-derived neuronal progenitors 48 h after stroke did not alter cortical infarct volume but preserved VPN neurons, reduced subsequent thalamic atrophy, and maintained diffusion properties of affected thalamocortical pathways.

These findings define secondary thalamic degeneration as a temporally ordered process in which early alterations in the thalamocortical pathway precede neuronal loss and delayed structural atrophy. Importantly, longitudinal tractography monitored both cortical stroke-induced thalamocortical degeneration and its modification by hiPSC-derived neuronal transplantation, establishing an in vivo approach to assess remote circuit degeneration and transplantation-dependent neuroprotection after cortical stroke.

**HIGHLIGHTS:**

- Cortical stroke causes progressive secondary thalamic degeneration
- Thalamic neuronal loss precedes delayed thalamic atrophy after stroke
- MR tractography reveals dynamic disruption of thalamocortical connectivity
- Human iPSC-derived progenitors protect thalamic neurons after stroke

## INTRODUCTION

Ischemic stroke is a leading cause of death and disability worldwide (Feigin, 2021), accounting for over 60% of all stroke cases (Pu et al., 2023). It results from interrupted cerebral blood flow, causing infarction, most commonly in the cerebral cortex. Cortical involvement occurs in about 60–70% of cases, with the middle cerebral artery (MCA) most frequently affected. Because the MCA supplies the primary motor and somatosensory cortices, 50–80% of stroke survivors develop sensory deficits, depending on lesion location (Kessner et al., 2019).

Stroke lesions often lead to secondary neurodegeneration in remote brain regions due to disruption of neural connectivity. In this context, cortical ischemia can induce secondary thalamic injury by disrupting reciprocal cortico-thalamic and thalamo-cortical connections. Clinical imaging studies have shown reduced cerebral blood flow, oxygen consumption, and glucose metabolism in the ipsilateral thalamus following cortical infarction (De Reuck et al., 1995; Nagasawa and Kogure, 1990; Ogawa et al., 1997; Reidler et al., 2018). MRI studies in patients with cerebral infarction sparing the thalamus have also identified alterations in thalamic signal intensity and diffusion properties on the infarct-ipsilateral side. More recently, Geng et al. provided MRI-based evidence for secondary remote thalamic atrophy after ischemic stroke and suggested that the extent of ipsilateral thalamic volume loss is associated with global post-stroke cognitive impairment (Geng et al., 2023). Supporting this, Johnston and collaborators, using a corticothalamic circuit model, found that chronic stroke patients showed reduced thalamic inhibition in the lesioned hemisphere, which was associated with secondary thalamic degeneration despite the absence of direct thalamic damage. This thalamic dysfunction was further associated with abnormal neural slowing and poorer cognitive and language outcomes (Johnston et al., 2024). Together, these observations indicate that cortical stroke can induce delayed abnormalities in remote thalamic structures and circuits. However, they do not establish whether early thalamic changes represent predominantly functional consequences of cortical disconnection or the onset of a progressive degenerative process leading to irreversible neuronal loss and structural atrophy.

Secondary thalamic injury has also been demonstrated in several experimental models of cortical stroke, including transient (Cao et al., 2017) and permanent (Iizuka et al., 1990) MCA occlusion and photothrombotic stroke (Aswendt et al., 2021). These studies have shown that the extent and distribution of secondary thalamic damage depend on the location and severity of the primary cortical lesion. Secondary neuronal damage is primarily attributed to retrograde degeneration, a process in which neuronal destruction spreads backward along axons after axonal injury (Fujie et al., 1990; Iizuka et al., 1990). Other mechanisms that may contribute to secondary thalamic injury include excitotoxicity (Paz et al., 2010; Ross and Ebner, 1990), apoptotic cell death (Fujie et al., 1990; Liang et al., 2007), amyloid-beta accumulation, blood-brain barrier (BBB) disruption (Yu et al., 2017), and neuroinflammation (Holmberg et al., 2009; Schroeter et al., 2006). Functional behavioral studies have further shown that reduced secondary thalamic damage can be associated with improved long-term behavioral performance after cortical stroke (Cao et al., 2017; Villa et al., 2007). Thus, secondary thalamic degeneration is an established consequence of cortical stroke and is potentially modifiable. Nevertheless, the existing literature has largely examined individual components of this process, such as neuronal degeneration, microglial activation, diffusion abnormalities, or delayed thalamic atrophy, at different time points and using different approaches. Consequently, the temporal relationship between these events remains incompletely defined.

It remains unclear how early disruption of thalamocortical pathways relates to later neuronal degeneration and tissue loss. Human MRI studies show diffusion abnormalities in the ipsilateral thalamus after MCA infarction, but their cellular basis and connection to irreversible degeneration are hard to determine. Experimental studies also show delayed thalamic neuronal loss and microglial activation, but the timeline between these responses and neuronal death isn’t fully understood in the same model. Early changes in thalamocortical connectivity may include axonal injury, disconnection, edema, and tissue responses, while later neuronal loss and volume reduction are more advanced and possibly irreversible. Clarifying this sequence is essential to understanding how cortical infarcts lead to remote degeneration and identifying windows for preserving vulnerable thalamic neurons.

A comprehensive longitudinal analysis of cellular and circuit-level measurements is needed to determine whether structural changes in thalamocortical pathways precede neuronal degeneration, whether neuronal loss occurs before thalamic atrophy, and how microglial activation timing relates to these events. This helps develop therapies targeting not just cortical infarcts but also remote neuronal networks connected to the injured cortex. Understanding these timing relationships doesn’t assume a single molecular cause but clarifies early circuit disruption versus later cellular degeneration and structural loss.

Several pharmacological approaches have been shown to reduce secondary thalamic degeneration after cortical stroke. These include inhibition of cathepsin B (Zuo et al., 2016), administration of osteopontin (Ladwig et al., 2019), suppression of caspase-1-mediated pyroptosis (Sun et al., 2026), pharmacological hypothermia induced by dihydrocapsaicin (Cao et al., 2017), and inhibition of autophagy through Beclin 1 knockdown or 3-MA treatment (Xing et al., 2012), among others. These interventions attenuated neuronal loss, neuroinflammation, and gliosis, while improving functional outcomes in experimental stroke models. Collectively, these findings indicate that secondary thalamic injury is not an inevitable consequence of cortical infarction but a therapeutically modifiable process. However, existing evidence has focused mainly on pharmacological or genetic manipulation of individual pathways, whereas less is known about whether restoring or reconstructing injured cortical circuitry can prevent degeneration in its remote thalamic targets.

Stem cell transplantation is a promising therapeutic strategy for stroke recovery because it facilitates cell replacement and promotes endogenous repair processes such as neuroplasticity, neurogenesis, and vascular remodeling (Palma-Tortosa et al., 2020). We have previously provided evidence that transplanted human induced pluripotent stem cell (hiPSC)-derived cortical neurons can improve behavioral recovery in the stroke-affected rodent brain through neuronal replacement and integration into injured host neural circuitry (Palma-Tortosa et al., 2020; Tornero et al., 2017). Recently, Li and co-workers (Li et al., 2023) reported that cortically grafted human embryonic stem cell-derived neural progenitors can alleviate secondary thalamic neurodegeneration after stroke in rats. However, whether hiPSC-derived cortical neurons, which can reconstruct components of injured cortical circuitry, can also preserve remote thalamic neurons and thalamocortical connectivity remains largely unexplored.

Based on these considerations, the present study had two complementary objectives. First, we aimed to define the temporal progression of secondary thalamic degeneration after cortical stroke, with particular emphasis on longitudinal diffusion MRI tractography to monitor changes in thalamocortical pathways. We combined repeated tractography and volumetric MRI with histological analyses of neuronal degeneration and microglial activation to relate early pathway disruption to subsequent neuronal loss and delayed thalamic atrophy. Second, we investigated whether early intracortical transplantation of cortically primed human hiPSC-derived neuronal progenitors could protect the remote thalamus and preserve thalamocortical connectivity. We used longitudinal tractography to assess both the progression of secondary pathway degeneration and its modification by cell transplantation.

Together, these experiments establish a temporal framework for secondary thalamic degeneration and evaluate tractography as an in vivo approach to monitor both remote neurodegeneration and transplantation-dependent neuroprotection after cortical stroke.

## METHODS

### Long-term neuroepithelial stem (lt-NES) cell line maintenance and cortical priming

Human iPS cell-derived lt-NES cells were generated from skin fibroblasts as previously described (Grønning Hansen et al., 2020) and expanded on plates coated with 0.1 mg/mL poly-L-ornithine and 10 mg/mL laminin (both from Sigma) in proliferation media (DMEM-F12 without HEPES + Glutamine, N2 at 1:100, and Glucose at 1.6 g/l), supplemented with bFGF (10 ng/mL, Peprotech), EGF (10 ng/mL, Peprotech) and B27 (1:1000, Invitrogen). The cells were passaged at a ratio of 1:3 every second to third day using trypsin (Sigma). For cortical priming, the growth factors (FGF, EGF) and B27 were omitted, and the cells were cultured at low density in differentiation-defined medium with bone morphogenetic protein 4 (BMP4) (10 ng/mL, R&D Systems), wingless-type MMTV integration site family, member 3A (Wnt3A) (10 ng/mL, R&D Systems), and cyclopamine (1 μM, Calbiochem) for 4 days. For a detailed protocol, see Palma-Tortosa et al., 2022(Palma-Tortosa et al., 2022).

### Animals

Twenty-seven Sprague-Dawley rats (251-275 g, used in sham and stroke-subjected groups) and 15 Nude Rats (Crl:NIH-Foxn1 RNU, 8 weeks old, used for transplantation group) were obtained from Charles River Laboratories. The animals were housed in individually ventilated cages under standard temperature and humidity conditions, along with a 12-hour light/dark cycle and unrestricted access to food and water. All procedures were conducted in accordance with the European Union Directive 2010/63/EU and had received approval from the ethical committee for laboratory animal use at Lund University and the Swedish Department of Agriculture.

### Distal middle cerebral artery occlusion

Focal ischemic injury was induced in 12- to 13-week-old rats by distal MCA occlusion (dMCAO) as described (Chen et al., 1986) with some modifications. Briefly, animals were anesthetized with isoflurane mixed with air, and the temporal bone was exposed. A 3 mm craniotomy was performed; the dura mater was carefully opened, and the cortical branch of the MCA was permanently ligated with a suture, cauterized, and cut. We isolated and ligated both common carotid arteries for 30 to 120 min. We closed the surgical wounds and allowed the animals to wake up during the ischemic period. Sham animals underwent the same surgery but without ligation of the MCA.

Animals with less than 80% reduction of ipsilateral cortex (quantified as Volume ipsilateral cortex/contralateral cortex*100) after dMCAO were excluded from the study.

### Intracortical transplantation

Intracortical transplantation of cortically fated lt-NES cell-derived progenitors, transduced with a lentivirus carrying a GFP reporter, was performed stereotaxically 48 h after dMCAO under anesthesia, as previously described (Palma-Tortosa et al., 2020). Before surgery, the cortically primed cells were resuspended on day 4 of differentiation in cytocentrifugation buffer to a final concentration of 100,000 cells/µL. A volume of 1 μL was injected into 2 deposits at the following coordinates (relative to bregma and brain surface): Site 1: Anterior/posterior: +1.5 mm, medial/lateral: 1.5 mm, dorsal/ventral: −2.0 mm; Site 2: Anterior/posterior: +0.5 mm, medial/lateral: 1.5 mm, dorsal/ventral: −2.5 mm.

### Tissue processing and immunohistochemistry

Animals were sacrificed at different time points after stroke surgery by injection of pentobarbital followed by transcardial perfusion of saline and 4% paraformaldehyde (PFA) solution. Brains were extracted and left in PFA for post-fixation for 24 h, then transferred to 20% sucrose solution until saturated. Brains were cut coronally (30 μm) on a microtome (Leica SM 2000 R). We stored sections in an anti-freezer solution until further use.

Free-floating coronal sections were preincubated in a blocking solution containing 5% normal donkey serum (NDS) and 0.25% Triton X-100 in 0.1 M phosphate-buffered saline (PBS).

For immunofluorescence, primary antibodies (chicken anti-GFP (Merck Millipore, 1:1000), rabbit anti-NeuN (Abcam, 1:500), and goat anti-Iba1 (Biolegend, 1:200)) were diluted in blocking solution and applied overnight at 4°C. The following day, sections were incubated with fluorophore-conjugated secondary antibodies (donkey anti-chicken A488, donkey anti-rabbit A488, or donkey anti-goat Cy3, all from Jackson Immunoresearch) for 2 h at room temperature. Nuclei were counterstained with DAPI (Sigma) for 10 min before mounting on gelatin-coated slides using Dabco (Sigma).

### Fluoro-Jade C staining

Sections were mounted on coated slides, and the staining protocol was followed according to the instructions of the Fluoro-Jade C staining kit (Biosensis). Briefly, tissue was treated with 9 parts 80% ethanol and 1 part Solution A (sodium hydroxide), submerged in a 50 mL tube for 5 min, then sequentially transferred to 70% ethanol for 2 min and distilled water for 2 min. Slides were incubated in 9 parts distilled water and 1 part Solution B (potassium permanganate) for 5 min, optimizing the initial 10 min incubation to reduce autofluorescence and enhance contrast.

To improve fluorescence intensity, prepare a staining solution of 7 parts distilled water, 2 parts Solution C (Fluoro-Jade C), and 1 part DAPI. Slides were incubated in the dark for 10 min, followed by three rinses in distilled water (1 min each) and dried at 50–60°C for 5 min. Slides were cleared in xylene for 3 min in the dark and coverslipped using a non-aqueous mounting medium such as DPX.

Fluoro-Jade C-labeled degenerating cells were visualized under blue light excitation, with an excitation peak of 485–506 nm and an emission peak of 525–527 nm.

### Imaging

Overview images of rat brain slices were taken using the extended focal imaging (EFI) brightfield or fluorescence mode at a 20X magnification in a Virtual Slide Scanning System (VS-120-S6-W, Olympus, Germany).

High-magnification images were taken using a confocal microscope (LSM780, Zeiss, Germany) and a 20X or a 40X immersion objective.

### Neuronal quantification

Infarct volume was quantified using NeuN-immunostained sections. The intact area, identified by NeuN+ cells in both the ipsilateral and contralateral hemispheres, was delineated and measured using ImageJ software. The infarct lesion was measured as cortical loss by dividing the healthy ipsilesional cortex (NeuN-stained cortical area) by the contralateral cortex. Data were expressed as a percentage (*100).

NeuN+ cell quantification was conducted using a deep learning-based application within Visiopharm software (version 3.5.1). The application identifies neurons stained with the NeuN antibody in the FITC channel within user-defined regions of interest (ROIs). It calculates both the quantity of NeuN+ cells and the total area within each ROI.

Using a trained model, AI-driven image analysis was applied to distinguish target cells from the background. Post-processing steps - including background removal, object dilation to resolve fragmentation, and object separation - were implemented to enhance accuracy. For each animal, ROIs were delineated to distinguish between the ipsilateral and contralateral regions.

### Microglial quantification

Total number of Iba1+ cells in the cortex was manually counted using ImageJ software to accurately assess microglial cell density in the peri-infarct regions. For that, Iba1+ cells were quantified in three anatomically distinct brain sections (1.70, −0.26, and −3.14 mm from bregma) to provide a rostro-caudal representation of the evolution of inflammation throughout the brain. Within each section, four standardized 500 × 500 μm square areas were selected adjacent to the infarct. These areas were positioned as follows: (*a*) above the infarct on the ipsilesional hemisphere, (*b*) below the infarct on the ipsilesional hemisphere, (*c*) contralateral to area (*a*), and (*d*) contralateral to area (*b*). This spatial sampling enabled a comparative analysis of microglial activation between peri-infarct tissue and corresponding contralateral sites. Means of (*a*, *b*) and (*c*, *d*) were calculated.

Because of the high density of microglial cells and overlapping processes in the thalamus after stroke, we could not quantify individual cells. For that reason, we calculated Iba1+ density in the ipsilateral and contralateral VPNs as the area covered by Iba1+ staining divided by the VPN area (area within the ROI). We determined this using a threshold-based area measurement protocol in Visiopharm software.

### Magnetic resonance imaging

Magnetic resonance imaging (MRI) to evaluate brain anatomy and fiber-tract connectivity was performed before dMCAO surgery and at 1–3 days, 1–2 weeks, and monthly for up to 3 months post-stroke. T2-weighted (T2w) anatomical imaging was used to assess morphology, and diffusion tensor imaging was used for tractography-based connectivity. Animals were initially anesthetized with O2 mixed with 3% isoflurane and then reduced to 1.8-2% for the duration of the imaging procedure. We monitored breathing and body temperature throughout the procedure. The procedure was performed using a 9.4 T preclinical MRI horizontal-bore scanner (Agilent, Santa Clara, USA) equipped with Bruker BioSpec AVIII electronics operating with ParaVision 7.0.0 and a BGA 12S HP gradient system (Bruker, Ettlingen, Germany) with a maximum gradient strength of 670 mT/m and a rise time of 130 ms. We used a quadrature volume resonator (112/087) for transmission and a rat brain phased-array coil for reception. All coils were from Bruker.

#### Anatomical imaging

We obtained high-resolution T2w images using a 2D rapid acquisition with relaxation enhancement (RARE) sequence. The following parameters were used: echo time (TE) 36 ms, RARE factor 8, spectral bandwidth 75 kHz, repetition time (TR) 3.3 s, resolution 90 x 90 mm^2^, field of view (FOV) 18 x 14 mm^2^, and slice thickness 0.8 mm. We used radiofrequency (RF) pulses with calculated sharpness 3 (Bruker terminology) and a bandwidth of 2 kHz, with a 2.1 ms excitation pulse and a 1.7 ms refocusing pulse. Thirty-four slices were acquired axially with 8 averages in 8 m 21 s.

#### Diffusion tensor imaging

The imaging slice positions and FOV for the DTI-EPI scans were taken from the anatomical imaging. We used TE 19 ms, spectral bandwidth 179 kHz, partial FT 1.6, TR 3 s, and resolution 280 x 280 mm^2^. We acquired 256 directions with b = 2000 s/mm^2^ using HARDI diffusion encoding, plus five additional images without diffusion encoding. RF pulses with calculated sharpness 4 (Bruker terminology) and a bandwidth of 2.2 kHz were used, with 2.7 ms pulse length for excitation and 2.3 ms for refocusing. We used one average, and the acquisition time was 13 m 3 s.

### Tractography

We further processed the T2w and DTI images with AIDAmri, which uses DSI Studio (Yeh, 2025) for deterministic fiber tracking. AIDAmri was previously described in detail for mouse brain MRI and registration with the Allen Mouse Brain Atlas (Pallast et al., 2019) and is available on the software homepage, including a user manual (https://github.com/Aswendt-Lab/AIDAmri). We modified the configurations to match the larger brain size of rats and to register with the SIGMA Rat Brain Atlas (Barriere et al., 2019). Briefly, pre-processing included converting raw Bruker images to NIfTI files in the BIDS structure (Gorgolewski et al., 2016). We bias-field-corrected T2w files and brain-extracted them using FSL BET. On the brain-extracted images, we delineated stroke masks with ITK-SNAP (Yushkevich et al., 2006) and imported them into AIDAmri. DTI data were processed with anisotropic diffusion filtering and slice-wise motion correction using FSL mcflirt to remove noise and motion before brain extraction using FSL bet. We applied two-step affine and non-linear registration of the SIGMA atlas to the T2w and DTI data. We corrected orientation by aligning the images with the standard anatomical atlas.

Whole-brain fiber tracking and quantitative scalar maps (fractional anisotropy; FA) were calculated using DSI Studio version 2023 (Yeh et al., 2013). We performed deterministic fiber tracking across the entire brain using the SIGMA template, which encompasses 234 segmented regions. The tracking process utilized parameters such as fiber length >0.4 or <24 mm, turning angle <55°, and tracking interval to ensure accurate reconstruction of brain fiber tracts.

### Statistical analysis

We performed statistical analyses using Prism 10 software (version 10.4.1, GraphPad). One-way ANOVA, followed by Tukey’s multiple-comparisons test was used to analyze a single variable across different time points. Two-way ANOVA followed by Sidak’s multiple-comparisons test was used to compare two variables (time and ischemia). We used simple linear regression to compare VPN versus cortical loss and VPN loss versus neuronal loss. Significance was set at p < 0.05. Data are presented as mean ± SD.

## RESULTS

### 1. Cortical ischemic stroke causes early and stable lesions in the somatosensory cortex followed by progressive secondary thalamic injury

To elucidate the timeline of lesion development in both the primary and secondary affected areas (cortex and thalamus, respectively), animals were sacrificed at different time points after stroke: 3-4 days, 2 weeks, 3, 4, and 6 months. Sham animals, subjected to surgery but without dMCAO, were used as controls.

We first quantified the stroke-induced cortical lesion in sections stained for the neuronal marker NeuN by measuring cortical tissue loss in the ipsilateral hemisphere relative to that on the contralateral side. We observed that the cortical damage was located in the primary somatosensory cortex barrel field (S1BF) and clearly detectable as early as 3-4 days after the insult, with no significant changes in tissue loss at later time points (**Figure 1A-B**).

**Figure 1.**
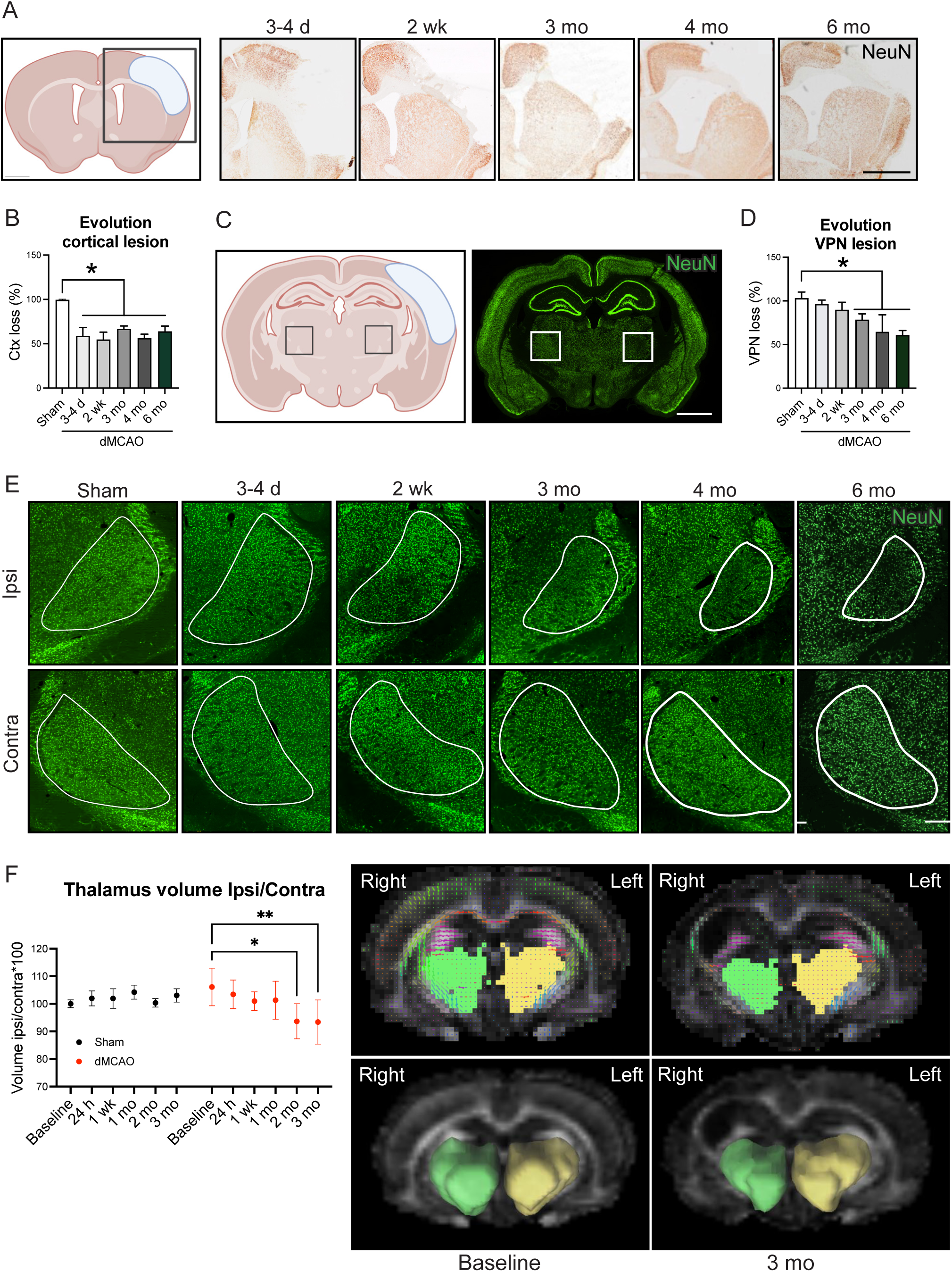
Temporal progression of the primary cortical and secondary thalamic lesions in a rat model of cortical ischemic stroke. (**A**) Schematic representation of the location of the cortical lesion following distal middle cerebral artery occlusion (dMCAO) in rats and representative brightfield images of the neuronal nuclei (NeuN)-DAB immunostaining in rat brain slices at different time points post-stroke. Scale bar, 2 mm. (**B**) Quantification of the cortical lesion at different time points post-stroke, measured as cortical loss in percentage (Area Ctx Ipsi/Area Ctx contra × 100). Ctx: cortex. *p<0.05. Data are presented as mean ± SD. (**C**) Schematic and overview of NeuN immunofluorescence in a caudal rat brain slice containing the ventral posterolateral nuclei (VPN). Scale bar, 2 mm. (**D**) Quantification of the percentage of reduction in VPN area calculated as (Area VPN ipsilateral/ Area VPN contralateral) × 100, at different time points post-stroke. *p<0.05. Data are presented as mean ± SD. (**E**) Representative immunofluorescence images of NeuN staining in rat brain slices at different time points post-stroke. Ipsilateral and contralateral VPN are outlined in white. Scale bar, 500 mm. (**F**) Relative volume [(ipsilateral/contralateral) × 100] of thalamus over time from magnetic resonance imaging, and representative images of 2D (upper) and 3D (lower) AIDAmri segmentations of bilateral thalamus at different time points post-stroke (Coronal view). Green, right thalamus; Yellow, left thalamus. *p<0.05 and **p<0.01. Data are presented as mean ± SD.

We then measured the size of the ipsilesional and contralesional VPN, a thalamic area with extensive projections to and from the somatosensory cortex, at multiple post-stroke time points. No differences were found in VPN area between the contralateral and ipsilateral hemispheres in sham animals. Also, the contralateral VPN area in stroke-subjected animals did not differ from that of their time-matched shams (**Supplementary Figure 1A,B**), indicating that the effect of cortical stroke on VPN is ipsilateral. Based on these findings, in all subsequent analyses, we used the contralateral VPN area as the control for each stroke-subjected rat, allowing us to account for variability among individual animals.

Three to four days and two weeks after the stroke, no changes in the ipsilateral VPN area of stroke animals were observed compared with sham animals. However, after 3 months, the ipsilateral VPN area decreased and then did not change through the latest time point examined at 6 months after stroke (**Figure 1C-E**). Notably, from 4 months onwards, the magnitude of thalamic damage, measured as the relative decrease in the ipsilateral VPN area, significantly correlated with that of the cortical lesion (Supplementary **Figure 1D**).

Consistent with these histological results, volume analysis of MRI images revealed a gradual decrease in relative (ipsilateral vs. contralateral) thalamic area, reaching a significant difference compared with sham at 2-3 months after stroke (**Figure 1F**).

### 2. Neuronal loss is the primary contributor to the thalamic damage

To further investigate the tissue damage in the cortex and VPN, we performed Fluoro-Jade C staining, a method sensitive to early signs of neuronal degeneration, at 2 time points: 2 weeks, when VPN tissue loss was not yet detectable in NeuN-stained sections, and 4 months post-stroke, when VPN atrophy was stable. No degeneration was detected in sham animals or in the contralateral hemisphere of stroke-affected rats (**Supplementary Figure 2A,B**). However, significant degeneration was observed in both the stroke-affected cortex and the ipsilateral VPN at both time points (**Supplementary Figure 2B**), indicating that thalamic degeneration begins before tissue loss is detectable.

We wanted to determine the role of neuronal loss for the secondary thalamic lesion after the cortical stroke. No significant changes in the number of NeuN+ neurons were observed in VPN of sham animals or in the contralateral hemisphere of stroke-affected rats (**Supplementary Figure 1C**). Therefore, we counted the numbers of NeuN+ cells in the ipsilateral VPN and compared them with those in the corresponding non-injured structures. Although no differences were observed 3-4 days after the stroke, significant neuronal loss in the ipsilateral VPN was detected already after 2 weeks and remained stable at later time points (**Figure 2A-C**). Importantly, the loss of neurons in the ipsilateral VPN correlated with a decrease in VPN size starting 3 months after stroke onset (**Figure 2D**). Moreover, we observed a positive correlation between cortical damage and loss of neurons in the ipsilateral VPN at 6 months post-stroke (**Supplementary Figure 3A-E**).

**Figure 2.**
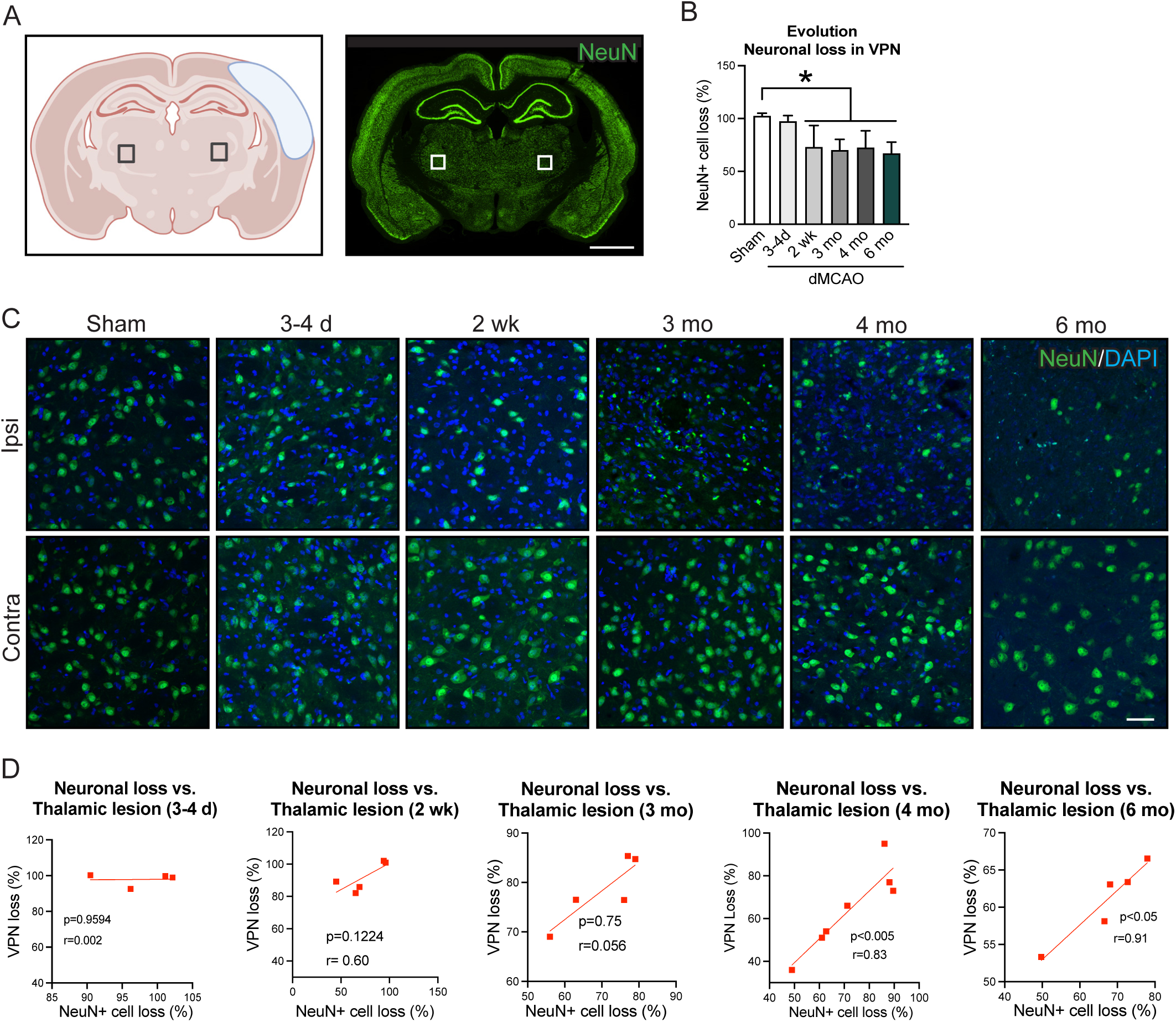
Progression of neuronal loss in the ventral posterior nuclei (VPN) following a cortical ischemic stroke in rats. (**A**) Schematic representation of the area where representative images for the neuronal nuclei (NeuN) were captured. Scale bar: 2 mm. (**B**) Quantification of the relative loss of NeuN+ cells ((NeuN+ cells in ipsilateral VPN / NeuN+ cells in contralateral VPN) × 100) in the VPN over time. *p<0.05. Data are presented as mean ± SD. (**C**) Representative immunofluorescence images of the NeuN+ cells in the ipsilateral and contralateral VPN at various time points after the stroke. Scale bar: 50 mm. (**D**) Linear regression analysis showing the correlation between the neuronal loss in the VPN and the VPN reduction at 3-4 days, 2 weeks, 3 months, 4 months, and 6 months post-stroke. N=4 −7 animals per time point.

These results provide evidence that (i) neuronal loss precedes detectable tissue atrophy in the ipsilateral VPN, (ii) VPN atrophy and neuronal loss is correlated with the extent of neuronal death in the cortex, which conceivably is the primary driver of thalamic degeneration after stroke.

### 3. Tractography shows axonal injury and demyelination in tracts connecting the ipsilateral thalamus and the stroke-affected cortex

Next, we used tractography to visualize axonal injury and demyelination in tracts connecting the ipsilateral thalamus with the damaged cortex (**Figure 3**). Tractography was performed using AIDAmri processing and DSI Studio on T2-weighted and DTI datasets. Fiber tracking was conducted between selected ROIs to assess stroke-related white-matter alterations. Four tractography-derived diffusion metrics were analyzed (**Supplementary Figure 4)**. Fractional anisotropy (FA), which reflects the directional coherence and structural integrity of axonal bundles, decreases when tracts are disrupted. Mean diffusivity (MD) represents the overall magnitude of water diffusion, with increases indicating reduced tissue density, edema, or necrosis. Axial diffusivity (AD) measures diffusion parallel to axonal fibers and decreases with axonal injury or degeneration. Radial diffusivity (RD) captures diffusion perpendicular to axons and increases when myelin integrity is compromised (Abdelrahman et al., 2022).

**Figure 3.**
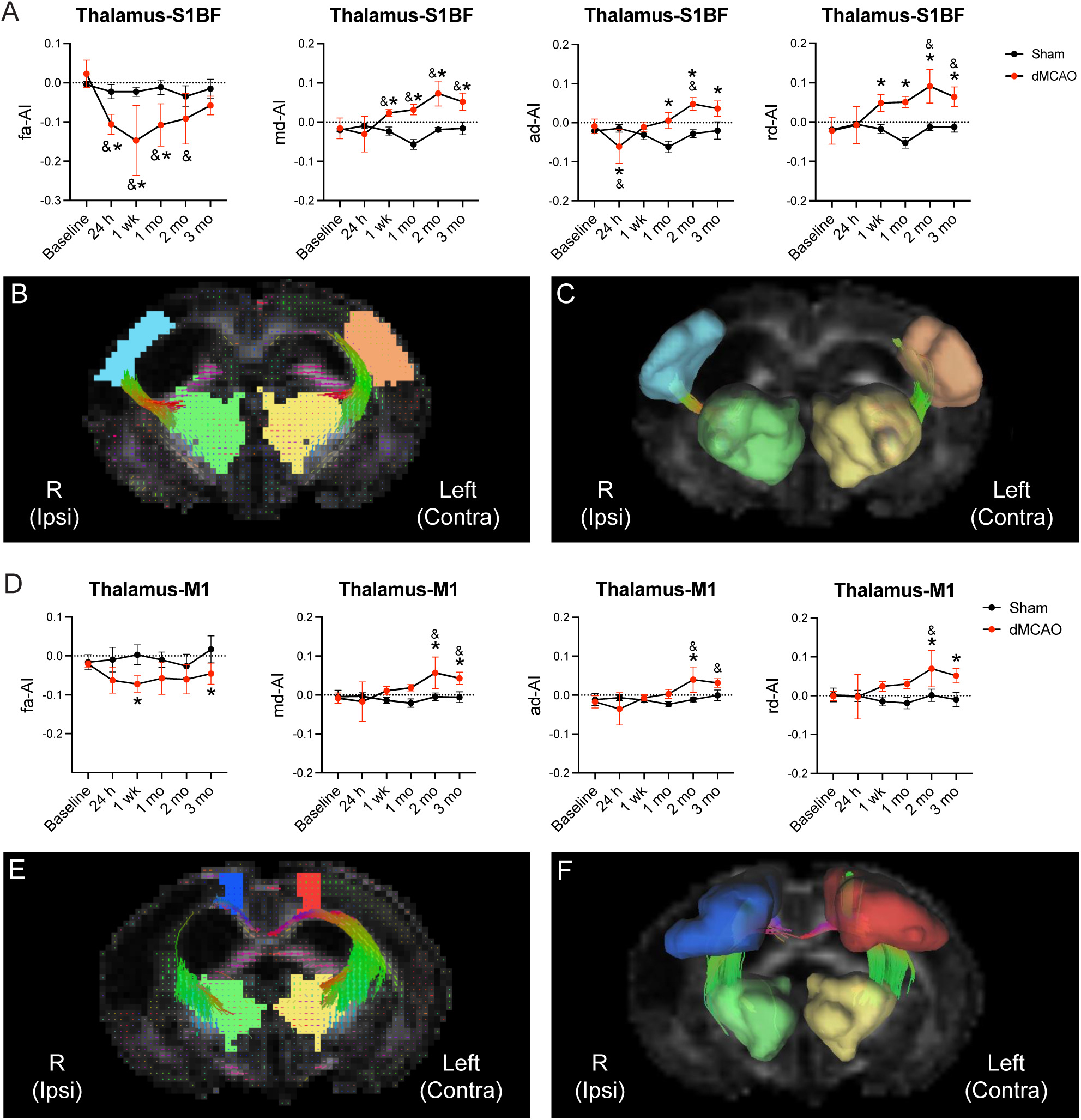
Tractography dynamics of the tracts connecting the ipsilateral thalamus and the stroke-affected cortex. (**A**) Asymmetry index (AI) of FA, MD, AD, and RD of thalamus-primary somatosensory cortex barrel field (S1BF) tracts at different time points post-stroke. \***p<0.05 and **p<0.01**. Data are presented as mean ± SD. (**B-C**) Representative (**B**) 2D and (**C**) 3D figures demonstrate the fiber tracts connecting the ipsilateral thalamus and S1BF (Coronal view). (**D**) AI of FA, MD, AD, and RD of thalamus-primary motor cortex (M1) tracts. (**E-F**) Representative (**E**) 2D and (**F**) 3D figures demonstrate the fiber tracts connecting the ipsilateral thalamus and S1BF (Coronal view) at different time points post-stroke. Green, right thalamus; Yellow, left thalamus; Sky blue, right S1BF; Orange, left S1BF; Blue, right M1; Red, left M1. Asymmetry index = (ipsilateral – contralateral)/ (ipsilateral + contralateral). N=4 animals per time point. *p<0.05 Sham vs. dMCAO, and while ^&^ p<0.05 dMCAO vs. Baseline. Data are presented as mean ± SD.

Tracts connecting the ipsilateral thalamus and the primary somatosensory barrel field (S1BF) showed a marked reduction in FA as early as 1 day post-stroke, reaching a minimum at 1 week and partially recovering thereafter. Consistent with tissue injury, MD and RD were significantly elevated compared to sham and baseline from 1 week onward, indicating edema and demyelination. AD decreased on day 1, consistent with acute axonal injury, and then gradually increased over time (**Figure 3A-C**). Tracts connecting the thalamus and the primary motor cortex exhibited similar temporal trends, though with smaller effect sizes (**Figure 3D-F**).

Together, these diffusion metrics reveal dynamic, time-dependent signatures of acute axonal injury, subsequent partial recovery, and evolving demyelination within stroke-affected thalamocortical pathways. In contrast, tracts linking the thalamus to regions largely spared by the primary lesion, such as the cingulate cortex (Cg1) (**Supplementary Figure 5A,C**) and the striatum (**Supplementary Figure 5B,D**), showed no significant differences between stroke and sham groups across all metrics.

### 4. Cortical ischemic stroke induces microglial activation in the cortex and ipsilateral VPN

We wanted to explore the possible interaction between activated microglia and neuronal loss in the somatosensory cortex, where the primary lesion occurs, and in the ipsilateral and contralateral VPNs (**Figures 4 and 5**).

**Figure 4.**
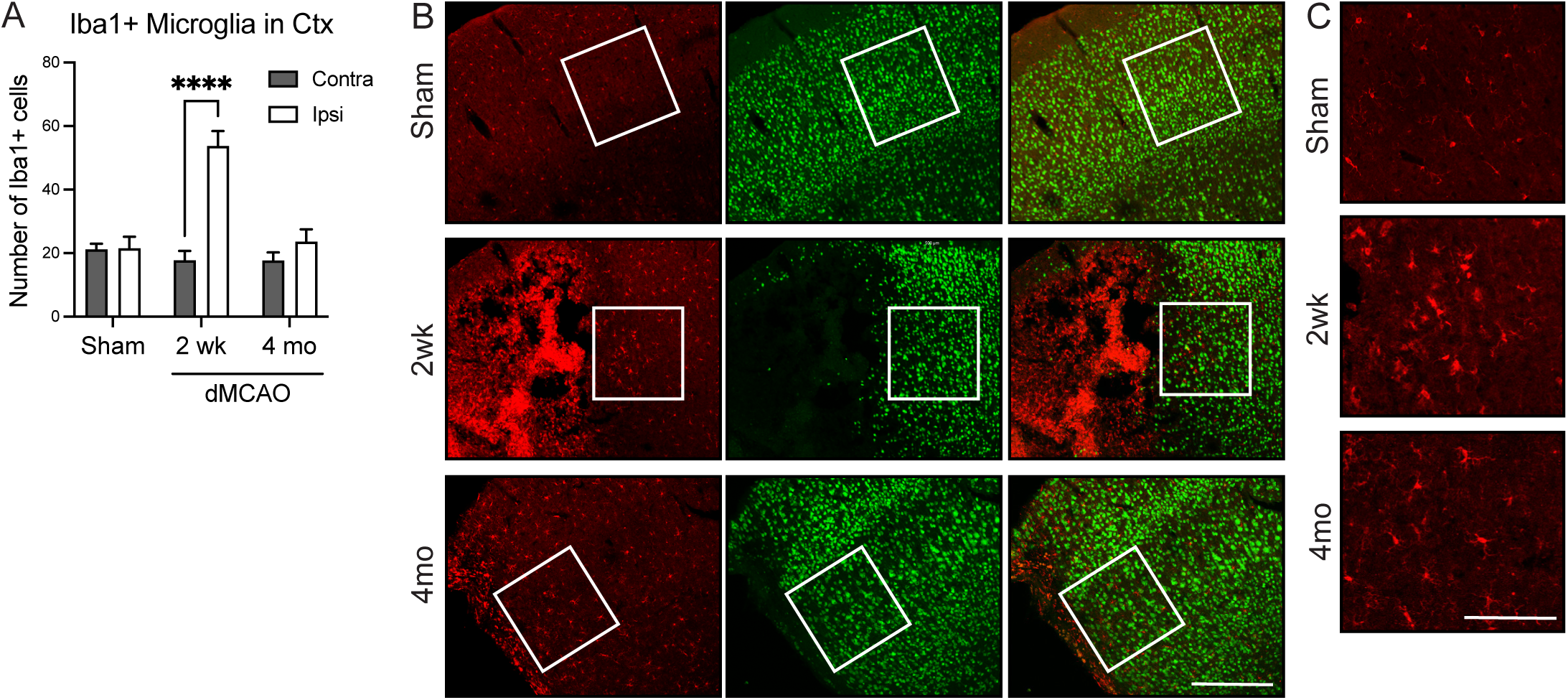
Time course of microglial activation in the somatosensory cortex after cortical ischemic stroke in rats. (**A**) Quantification of Iba1+ cells in the peri-infarct area and the corresponding area in the contralateral hemisphere in sham animals and in animals at 2 weeks and 4 months after stroke. N = 5–6 animals per time point. *p<0.05. Data are presented as mean ± SD. (**B**) Representative images of the microglial marker Iba1 and the neuronal marker NeuN, showing the area where Iba1+ cells were quantified in sham animals and at 2 weeks and 4 months post-stroke. Scale bar, 500 mm. (**C**) High-magnification representative immunofluorescence images of the areas where Iba1+ cells were quantified. Scale bar, 500 μm.

**Figure 5.**
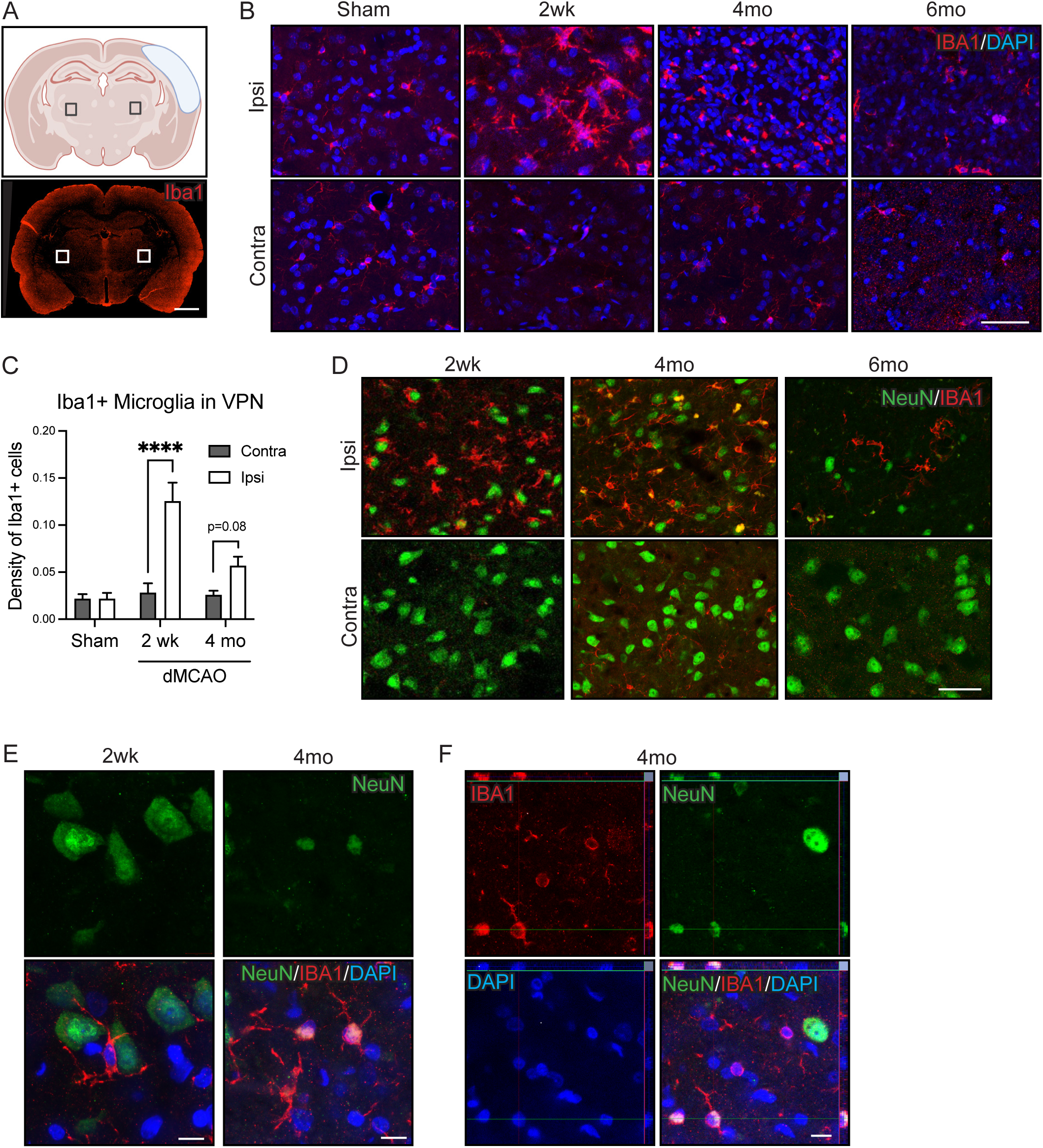
Time course of microglial activation in the ipsilateral ventral posterior nuclei (VPN) of a rat model of cortical ischemic stroke. (**A**) Schematic representation indicating where representative images of the microglial marker Iba1 were taken. Scale bar, 2 mm. (**B**) Representative immunofluorescence images showing the microglial marker Iba1 and the nuclear marker DAPI in the ipsilateral and contralateral VPN at various time points post-stroke. (**C**) Quantification of the Iba+ density (Iba1+ area/VPN area) at 2 weeks and 4 months post-stroke. ****, p<0.001. Data are presented as mean ± SD. (**D**) Representative immunofluorescence images displaying Iba1 and neuronal nuclei (NeuN) staining in the ipsilateral and contralateral VPN at different time points post-stroke. Scale bar, 50 mm. (**E**) High-magnification images of Iba1- and NeuN-immunopositive cells at 2 weeks post-stroke, where both markers were in close proximity, and at 4 months post-stroke, where colocalizations were observed. Scale bar, 20 mm. (**F**) Confocal orthogonal views illustrating the colocalization of NeuN and Iba1 at 4 months post-stroke. Scale bar, 20 mm.

The number of Iba1+ microglial cells increased 2 weeks post-stroke in the ipsilateral cortex compared to contralateral and sham animals. At 4 months, microglial number had returned to basal levels (**Figure 4A-C**).

In the thalamus, the area covered by Iba1+ staining in the ipsilateral VPN increased at 2 weeks post-stroke. By 4 months, although the Iba1+ area had declined, it still showed a trend toward higher levels compared with the contralateral side (**Figure 5B-C**). By 6 months post-stroke, microglial density levels were comparable to those observed in sham animals (**Figure 5C**). Qualitative analysis showed that at 2 weeks post-stroke, Iba1+ microglia were frequently located near intact NeuN+ neurons, suggesting early microglia-neuron interactions. At 4 months, colocalization of Iba1 and NeuN staining within the same cell became evident, potentially indicating ongoing phagocytosis of damaged neurons (**Figure 5D-F**). In contrast, at 6 months post-stroke, no colocalization between NeuN and Iba1 was detected (**Figure 5D**), which might reflect the resolution of the inflammatory response and the complete clearance of damaged neurons. No changes in the number or appearance of Iba1+ cells were observed in the contralateral VPN at any time point (**Figure 5B**).

### 5. Intracortical transplantation of iPS cell-derived progenitors rescues the stroke-induced neuronal loss in VPN

To transplant human-derived cells into rat brains (xenotransplantation) and avoid immune rejection, we used nude athymic rats for dMCAO induction. Forty-eight hours later, cortically primed human lt-NES cell-derived progenitors were transplanted in close proximity to the cortical injury. After six months, we sacrificed animals and processed them for immunohistochemistry.

We first assessed whether intracortical transplantation of human lt-NES cell-derived progenitors affected the extent of the cortical damage. We observed no differences in cortical lesion size between transplanted and non-transplanted animals (**Figure 6A**), consistent with our previous findings (Tornero et al., 2017).

**Figure 6.**
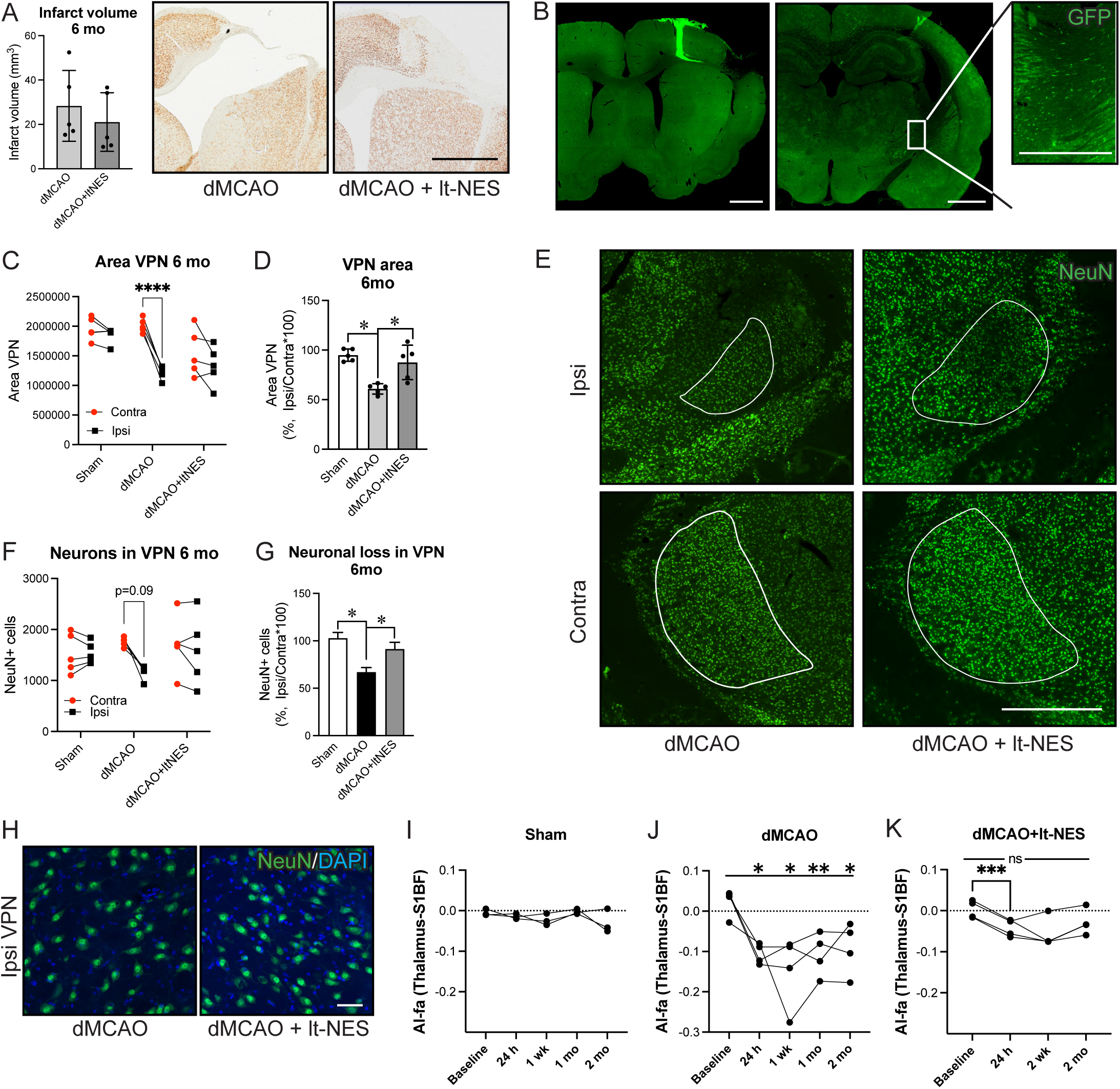
Early intracortical transplantation of human long-term neuroepithelial stem (lt-NES) cell-derived progenitors inhibits secondary neuronal degeneration in the ipsilateral ventral posterior nuclei (VPN) of the thalamus in a rat model of cortical ischemic stroke. (**A**) Quantification of infarct volume 6 months post-stroke in transplanted and non-transplanted rats. Representative brightfield images of NeuN-DAB immunostaining in rat brain slices. Scale bar, 2 mm. (**B**) Overview of graft location in rostral and caudal brain slices. Lt-NES cells are labeled with GFP for visualization. Scale bar: 2 mm (low magnification), 500 μm (high magnification). (**C**) Quantification of ipsilateral and contralateral VPN area in sham, stroke, and transplanted rats. (**D**) The percentage of reduction in VPN area is calculated as (Ipsilateral VPN Area / Contralateral VPN Area) × 100. (**E**) Representative immunofluorescence images of NeuN staining in brain slices of stroke-subjected (dMCAO) and stroke-transplanted (dMCAO + ltNES) rats. Ipsilateral (Ipsi) and contralateral (Contra) VPN are outlined in white. Scale bar, 500 mm. (**F**) Quantification of NeuN+ cells in the ipsilateral and contralateral VPN. (**G**) Quantification of the relative loss of NeuN+ cells ((NeuN+ cells in ipsilateral VPN / NeuN+ cells in contralateral VPN) × 100) in the VPN. (**H**) High-magnification immunofluorescence images of NeuN staining in stroke-affected rat brain slices with and without transplantation. Scale bar, 50 mm. (**I**,**J**) Asymmetry index (AI) of FA of (**I**) thalamus-primary somatosensory cortex barrel field (S1BF) tracts and (**J**) thalamus-primary motor cortex (M1) tracts at 3 months post-transplantation. Asymmetry index = (ipsilateral – contralateral)/ (ipsilateral + contralateral). *p<0.05 and ****, p<0.001. Data are shown as mean ± SD.

We then examined the effect of intracortical transplantation on the stroke-induced secondary thalamic lesion. Six months after transplantation, graft location analysis showed that most grafted cells remained near the transplantation site in the frontal somatosensory cortex, while some graft-derived processes extended into the caudal part of the brain, especially the thalamus and internal capsule (**Figure 6B**). Notably, transplanted animals showed a similar ipsilateral VPN to the sham group and a significantly larger ipsilateral VPN than non-grafted stroke-lesioned animals (**Figure 6C-E**). Importantly, we observed no significant difference between the ipsilateral VPN of grafted animals and the corresponding area in sham animals.

We also investigated whether preserved VPN size after lt-NES cell transplantation was associated with increased neuronal survival in this region. We quantified the number of NeuN+ neurons in the ipsilateral VPN of transplanted and non-transplanted stroke-lesioned animals and observed a higher number of NeuN+ cells in the VPN of the transplanted group (**Figure 6F-H**).

Finally, we analyzed the effects of lt-NES cell-derived progenitor transplantation on thalamo-cortical tractography. While no changes were found in sham animals, the FA values of tracts connecting the ipsilateral thalamus and primary somatosensory cortex were affected by cortical stroke at all the different times studied (24 hours, 1 week, 1 month and 2 months). Interestingly, after an initial decrease 24 hours after stroke, consistent with values in the dMCAO group, intracortical transplantation of lt-NES cell-derived progenitors 48 hours after stroke surgery restored FA values at 2 weeks and remained similar to those corresponding to Sham animals until 2 months, the last time point studied. (**Figure 6I,K**). These results suggest that the cell grafts improved the tract diffusion directionality, possibly through sparing surviving axons or facilitating tracts regeneration, explanation supported by our histologic findings.

Taken together, lt-NES cell-derived transplantation mitigated secondary injury in the thalamus following cortical stroke by preserving VPN volume, increasing the number of neurons, and promoting tract connectivity between the thalamus and the cortex.

## DISCUSSION

This study used longitudinal diffusion MRI tractography to monitor stroke-induced degeneration of thalamocortical pathways and assess if transplantation of human iPSC-derived neuronal progenitors could modify these changes. Our objectives were to define the progression of secondary thalamic degeneration and to see if early intracortical transplantation could protect the thalamus and preserve thalamocortical connectivity. By combining longitudinal tractography, MRI, and histological analyses, we identified a sequence linking early pathway disruption to later neuronal loss and thalamic atrophy. The primary cortical infarct appeared within days, while diffusion changes in pathways were detected early. Neuronal degeneration in the VPN was evident at 2 weeks, with thalamic atrophy developing months later. Our findings show that longitudinal tractography tracks degeneration progression before overt thalamic loss. We also found that intracortical transplantation of human neuronal progenitors 48 h post-stroke preserved VPN neurons, reduced atrophy, and maintained pathway diffusion properties. This approach allows *in vivo* monitoring of neuroprotection. Because we did not assess functional outcomes, these findings do not prove that preserving the thalamus improves recovery. Although our previous work showed improved performance at 6 months, further study is needed to determine whether protection against secondary degeneration aids recovery.

Secondary thalamic abnormalities after cortical stroke have been reported in both experimental models and human patients, including changes in metabolism, diffusion, MRI signal, and ultimately thalamic volume. Human longitudinal MRI studies show that ipsilateral thalamic diffusion abnormalities emerge weeks to months after MCA infarction, consistent with progressive remote tissue injury rather than an acute primary thalamic lesion (Herve et al., 2005; Ogawa et al., 1997; Pappata et al., 2000). However, our combination of thalamocortical tractography with parallel histological time courses and repeated post-stroke measurements provides a more direct circuit-level description than measuring thalamic diffusion properties alone.

More recent studies have demonstrated chronic ipsilateral thalamic atrophy after non-thalamic infarction and linked the magnitude of atrophy to infarct size and cognitive impairment (Geng et al., 2023). However, these observations do not establish that all early thalamic abnormalities represent irreversible neuronal degeneration. Indeed, an important unresolved issue is whether early thalamic changes reflect transient functional alterations caused by loss of cortical input, reversible cellular or metabolic responses, or the initiation of a progressive degenerative process. Our findings help address this distinction by demonstrating that early alterations in thalamocortical pathways are followed by histologically detectable neuronal degeneration before macroscopic thalamic atrophy becomes apparent. The data therefore support a model in which functional and microstructural disconnection precedes and potentially contributes to subsequent neuronal loss, rather than matching early thalamic dysfunction with established structural degeneration.

A major finding of the present study is that secondary thalamic degeneration follows a delayed, regionally selective trajectory. The cortical infarct was clearly established by 3-4 days and remained relatively stable thereafter, whereas measurable reduction in ipsilateral VPN volume was not detected until approximately 3 months after stroke and then remained stable through 6 months. This temporal dissociation indicates that delayed VPN atrophy is not simply a passive extension of the primary cortical lesion. Instead, progressive loss of thalamic tissue occurs after a prolonged interval during which cellular and connectivity changes are already underway. Importantly, NeuN analysis demonstrated that significant neuronal loss in the VPN was detectable as early as 2 weeks, when no significant reduction in VPN volume was yet apparent. Fluoro-Jade C staining further revealed degenerating neurons in the ipsilateral VPN at this stage. The subsequent correlation between neuronal loss and VPN volume reduction suggests that neuronal death is a major cellular substrate of the delayed thalamic atrophy. Moreover, the correlation between cortical lesion size and both VPN neuronal loss and later VPN atrophy supports a circuit-dependent relationship between the severity of the primary cortical injury and the extent of remote thalamic degeneration. This extends previous observations that the magnitude and distribution of secondary thalamic injury depend on the cortical territory affected and its thalamocortical connections (Schroeter et al., 2006; Seitz et al., 1999).

Longitudinal tractography data add another dimension to this model by showing that structural alterations in thalamocortical pathways emerge well before detectable thalamic atrophy. In tracts connecting the ipsilateral VPN with the stroke-affected S1BF, FA decreased as early as 1 day after stroke and reached a minimum at approximately 1 week. At the same time, MD and RD increased from 1 week onward, whereas AD initially decreased and subsequently increased. Although diffusion metrics are not specific to individual pathological processes, their temporal sequence is consistent with acute axonal injury followed by evolving changes in tissue organization and myelin integrity. The partial recovery of FA after its early decline does not necessarily indicate complete structural recovery. Rather, it may reflect dynamic shifts in the relative contributions of axonal injury, edema, cellular responses, and subsequent tissue remodeling. Importantly, these abnormalities were selective for pathways connecting the thalamus with stroke-affected cortical regions, whereas pathways linking the thalamus with largely spared regions, such as the cingulate cortex and striatum, did not show comparable changes. This anatomical specificity strongly supports the interpretation that the observed alterations reflect disruption of the affected thalamocortical network rather than a generalized consequence of stroke.

Our findings therefore support a more integrated model of secondary thalamic degeneration. The primary cortical infarct immediately disrupts thalamocortical and corticothalamic axons, reflected in early changes in AD and FA. This acute structural disconnection is followed by progressive alterations in diffusion properties, consistent with evolving axonal and myelin pathology. At the cellular level, neuronal degeneration in the VPN becomes detectable by 2 weeks, before sufficient neuronal loss has accumulated to produce measurable structural atrophy. Over the following months, the cumulative loss of thalamic neurons and associated tissue results in delayed VPN shrinkage. The strong relationship between cortical lesion severity and subsequent VPN neuronal loss further suggests that the magnitude of the initial cortical injury determines the degree of remote circuit disruption. This model is consistent with previous human studies showing delayed increases in thalamic MD after MCA occlusion-induced stroke and with evidence that remote white-matter degeneration can progress for months after stroke (Liang et al., 2007; Pappata et al., 2000). However, the present study advances these observations by linking the temporal evolution of tract-level diffusion abnormalities directly to neuronal degeneration and subsequent anatomical atrophy within a defined thalamocortical circuit.

Post-stroke secondary degeneration of neural tracts, including the cortico-spinal tracts (Liang et al., 2007; Moller et al., 2007), cortico-cerebellar tract (Schulz et al., 2017), cortico-rubro-spinal tracts (Ruber et al., 2012), and arcuate fasciculus (Narimani et al., 2026), has been demonstrated using clinical DTI, which revealed disrupted axonal integrity and Wallerian degeneration *in vivo*. This structural connectivity plays an important role in functional outcomes (Koch et al., 2016), and diffusion tensor tractography metrics have proven valuable functional predictors after stroke, including motor (Kim et al., 2015) and speech domains (Narimani et al., 2026).

To our knowledge, previous studies have not longitudinally combined tractography of anatomically defined thalamocortical pathways with parallel histological assessment of neuronal degeneration and quantitative measurements of delayed thalamic atrophy after cortical stroke. Although longitudinal diffusion MRI studies have demonstrated progressive remote thalamic and other structural changes after stroke (Buffon et al., 2005; Herve et al., 2005), and experimental studies have combined longitudinal in vivo fiber tracking with histological validation of secondary thalamic neurodegeneration (Aswendt et al., 2021), our multimodal approach integrates longitudinal tractography of defined thalamocortical pathways with histological assessment of neuronal degeneration and quantitative thalamic volumetry to relate early pathway abnormalities to subsequent neuronal loss and structural thalamic degeneration.

The microglial response further supports the view that secondary thalamic injury is an active, temporally regulated process rather than delayed tissue loss from disconnection. In both the cortex and the ipsilateral VPN, microglial activation increased at approximately 2 weeks after stroke, coinciding with the period when neuronal degeneration was already detectable. However, the subsequent events varied between the primary and secondary lesions. Cortical microglial numbers returned toward basal levels by 4 months, whereas increased Iba1+ staining in the VPN persisted longer and only approached sham levels by 6 months. This prolonged response in the remote thalamus is consistent with secondary inflammation, in which injury-induced immune responses propagate along functionally connected neural networks beyond the primary lesion (Holmberg et al., 2009; Jones et al., 2018). Our observations add an important temporal component to this model: microglial activation in the VPN begins during early neuronal degeneration and persists through the transition from cellular degeneration to structural atrophy.

The spatial relationship between microglia and neurons also suggests a possible sequence of microglia-neuron interactions. At 2 weeks, Iba1+ microglia were frequently adjacent to apparently intact NeuN+ neurons, raising the possibility that microglial activation precedes or accompanies neuronal dysfunction rather than being restricted to clearing already-dead cells. At later time points, Iba1 and NeuN colocalization became evident, consistent with microglial engulfment of neuronal material during established neuronal degeneration. By 6 months, this colocalization was no longer observed, coinciding with normalization of microglial density and suggesting resolution of the active clearance phase. These observations are consistent with a model in which microglia initially respond to signals generated by remote circuit injury and then remove degenerating neuronal material. However, because we used Iba1 staining and qualitative assessment of Iba1/NeuN colocalization, we cannot establish whether microglial activation causally drives neuronal death or predominantly reflects a response to neuronal injury. The data nevertheless identify a temporally defined period in which microglia-neuron interactions coincide with the onset and progression of secondary thalamic degeneration. This interpretation is supported by studies showing that modulation of inflammatory pathways can reduce secondary thalamic neuronal loss and associated neurological deficits after cortical stroke (Liang et al., 2025).

The transplantation experiments provide evidence that this delayed degeneration is not an inevitable consequence of cortical stroke. Early intracortical transplantation of cortically primed human iPSC-derived neuronal progenitors did not alter the size of the primary cortical infarct but markedly reduced subsequent VPN atrophy and preserved NeuN+ neurons 6 months after stroke. Thus, the protective effect occurred remotely from the primary lesion and cannot be explained simply by reducing the initial cortical infarct. The finding that most grafted cells remained near the cortical transplantation site, while graft-derived processes extended toward the internal capsule and thalamus, provides a potential anatomical basis for this remote effect. Our previous studies demonstrated that human lt-NES cell-derived progenitors transplanted into the stroke-damaged cortex receive direct synaptic input from host thalamic neurons and can form synaptically functional thalamocortical connections (Tornero et al., 2017). In the present study, graft-derived processes extending toward the thalamus suggest that the graft may become incorporated into the damaged thalamocortical network. Nevertheless, because we did not directly assess functional connectivity, these results should be interpreted as anatomical evidence of graft-host integration rather than direct evidence of restored circuit function.

Several mechanisms may account for the preservation of thalamic neurons after transplantation. One possibility is that graft-derived cortical neurons reconstruct or partially replace disrupted cortical targets, thereby reducing the consequences of cortical denervation for connected thalamic neurons. The delayed onset of measurable VPN atrophy provides a potentially important therapeutic window during which transplanted neurons can mature and establish host-graft interactions before substantial remote tissue loss occurs. A second possibility is that grafted neurons exert trophic or paracrine effects that enhance the survival of vulnerable host neurons. A third possibility is modulation of the inflammatory environment. Although we did not directly measure cytokine production or microglial responses in transplanted animals, the prolonged microglial response observed in the untreated stroke group identifies neuroinflammation as a plausible component of the secondary injury process. Previous work has shown that transplanted neural progenitors can modify the post-stroke inflammatory environment, and Li et al. (Li et al., 2023) reported reduced secondary thalamic damage after transplantation of human embryonic stem cell-derived neural progenitors, potentially involving BDNF/TrkB signaling. Our findings extend these data by showing preservation of remote thalamic neurons over 6 months following transplantation of cortically specified human iPSC-derived progenitors.

The present study also extends previous grafting studies showing that cortical transplantation is associated with reduced thalamic atrophy after cortical injury. Fetal cortical grafts have been shown to establish connections with the thalamus and attenuate secondary thalamic atrophy after cortical lesions (Mattsson et al., 1997; Sorensen et al., 1996). Cortical grafting after neonatal cortical ablation has also been reported to transiently or permanently reduce thalamic atrophy (Haun and Cunningham, 1984; Sharp and Gonzalez, 1986). Our study demonstrates that a similar principle can be achieved using human iPSC-derived, cortically specified neuronal progenitors in an adult ischemic stroke model. Importantly, these data place this protective effect within a broader temporal framework: thalamic neuronal degeneration begins weeks before detectable tissue atrophy, while tract-level abnormalities are evident within days. This identifies a potential therapeutic window in which transplantation aiming for neuronal circuit reconstruction may preserve remote neuronal populations before irreversible structural degeneration develops.

These findings have implications for interpreting and monitoring secondary neurodegeneration after stroke. The delayed development of VPN atrophy means that conventional volumetric MRI may underestimate the extent of remote injury in the early and subacute stages. In contrast, diffusion-based measurements of thalamocortical pathways detected abnormalities shortly after stroke, preceding detectable thalamic volume loss.

Combining tractography with anatomical MRI and histological markers therefore provides complementary measures across different stages of the degenerative process. This is consistent with clinical studies showing that DTI can detect remote structural abnormalities in thalamic tissue and white-matter pathways after stroke, before or independently of overt structural atrophy (Liang et al., 2007; Pappata et al., 2000). Our findings further suggest that pathway-specific diffusion measures may serve as sensitive biomarkers for monitoring remote circuit injury and evaluating interventions designed to preserve connectivity.

Preventing secondary thalamic injury may be clinically relevant because the thalamus is a major integrative relay station for sensory, motor, cognitive, and cortical network functions. Human studies have linked secondary thalamic atrophy following non-thalamic infarction to poorer cognitive performance (Geng et al., 2023), while diffusion abnormalities in non-lesioned thalamic nuclei and their cortical projections have been associated with functional deficits after stroke (Keser et al., 2021). However, the present study does not establish that preserving VPN neurons or thalamocortical diffusion properties translates into improved behavioral recovery. Instead, it shows that remote thalamic neurodegeneration is a measurable and modifiable component of the post-stroke response and provides a mechanistic rationale for testing whether preserving this circuit improves long-term neurological function.

Several limitations should be acknowledged. First, we did not assess functional recovery, preventing a direct link between preserved thalamic integrity and connectivity and behavioral outcome. Future studies should therefore combine longitudinal behavioral testing with MRI and histological analyses to determine whether early preservation of thalamocortical pathways predicts improved functional recovery. Second, although the temporal association between microglial activation and neuronal degeneration suggests a potential role for neuroinflammation, the present experiments do not establish a causal relationship. Determining whether microglia drive, facilitate, or primarily respond to secondary neuronal degeneration will require selective manipulation of microglial activity or inflammatory signaling. Finally, diffusion metrics are sensitive to multiple biological processes and cannot independently distinguish axonal degeneration from demyelination, edema, or other changes in tissue organization. The strength of the present approach therefore lies in merging tractography with histological evidence rather than interpreting any individual diffusion metric in isolation. Future studies combining longitudinal behavioral assessment, cell-specific tracing, microglial characterization, and single-cell or spatial transcriptomics will be important for defining how transplanted neurons interact with host immune and neuronal populations to preserve remote neuronal circuits.

In conclusion, our study shows that longitudinal diffusion MRI tractography can monitor cortical stroke-induced degeneration of thalamocortical pathways and its modification by human iPSC-derived neuronal transplantation. By integrating tractography with MRI and histological analyses, we define secondary thalamic degeneration as a progressive process and demonstrate that early neuronal transplantation preserves both remote thalamic neurons and thalamocortical structural connectivity. These findings establish longitudinal tractography as a valuable *in vivo* approach for monitoring remote circuit degeneration and transplantation-dependent neuroprotection after cortical stroke.

## ACKNOWLEDGEMENTS

This work is supported by grants from the Swedish Research Council, Swedish Brain Foundation, Swedish Stroke Foundation, Regional Research Support from the Southern Swedish Healthcare Region, and the Swedish Government Initiative for Strategic Research Areas (StemTherapy). We thank the Lund University Bioimaging Center (LBIC) for providing experimental MRI resources.

## AUTHOR CONTRIBUTIONS

**Sopiko Kartsivadze**: investigation, methodology, formal analysis, validation. Writing - Review & Editing. **Yu-Ping Shen**: investigation, formal analysis, software. Writing - Review & Editing. **Linda Jansson**: investigation, Writing - Review & Editing. **Lucía Galán-Pintado**: investigation, Writing - Review & Editing. **Keshav Kher**: investigation, Writing - Review & Editing. **Sara Avilés-Carpintero**: investigation, Writing - Review & Editing. **Michael Gottschalk**: methodology, formal analysis, software, Writing – Review & Editing. **Aref Kalantari**: formal analysis, software, Writing - Review & Editing. **Daniel Mertens**: formal analysis, software, Writing - Review & Editing. **Raquel Martinez-Curiel**: investigation, Writing - Review & Editing. **Markus Aswendt**: software, Writing - Review & Editing. **Olle Lindvall**: writing-Original draft, Writing - Review & Editing. **Sara Palma-Tortosa**: conceptualization, writing-Original draft, supervision, funding acquisition. Writing - Review & Editing. **Zaal Kokaia**: conceptualization, writing-Original draft, project administration, funding acquisition, Writing - Review & Editing.

## SUPPLEMENTARY FIGURES

**Supplementary Figure S1.**
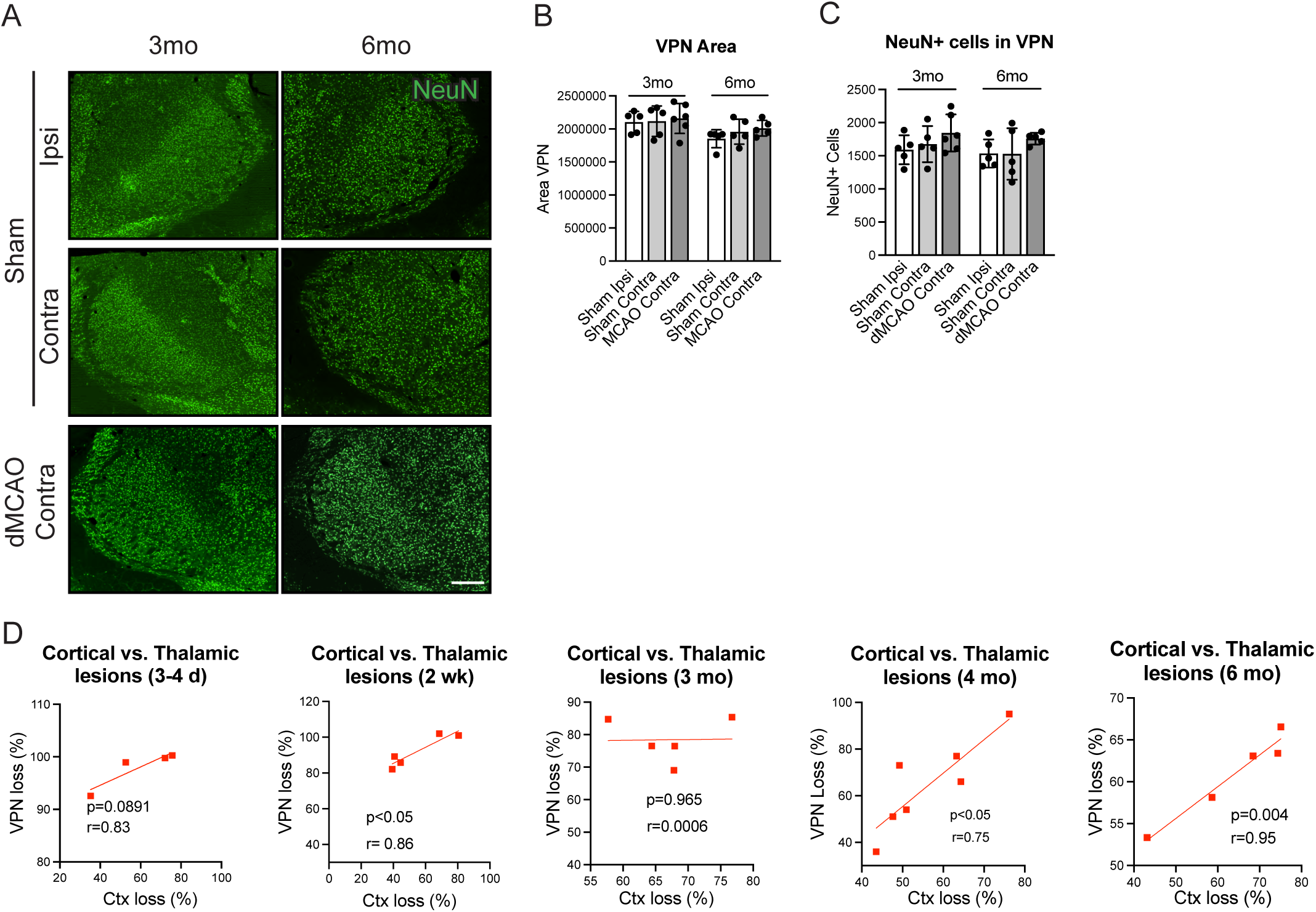
Assessment of the ventral posterior nuclei (VPN) area and number of neurons in the VPN of sham and non-lesioned hemisphere of stroke-affected animals. (**A**) Representative immunofluorescence images of NeuN-stained ipsilateral and contralateral VPN of sham-treated rats and rats at 3- and 6-months post-stroke. Scale bar, 500 mm. (**B-C**) Quantification of the VPN area (**B**) and NeuN+ cells (**C**) in the ipsilateral and contralateral VPN of sham-treated animals and contralateral VPN of stroke animals at 3- and 6-months post-stroke. (**D**) Linear regression analysis showing the correlation between cortical and VPN loss at 3-4 days, 2 weeks, 3 months, 4 months, and 6 months post-stroke. N = 4 - 7 animals per time point. *p<0.05. Data are shown as mean ± SD. Data are shown as mean ± SD.

**Supplementary Figure S2.**
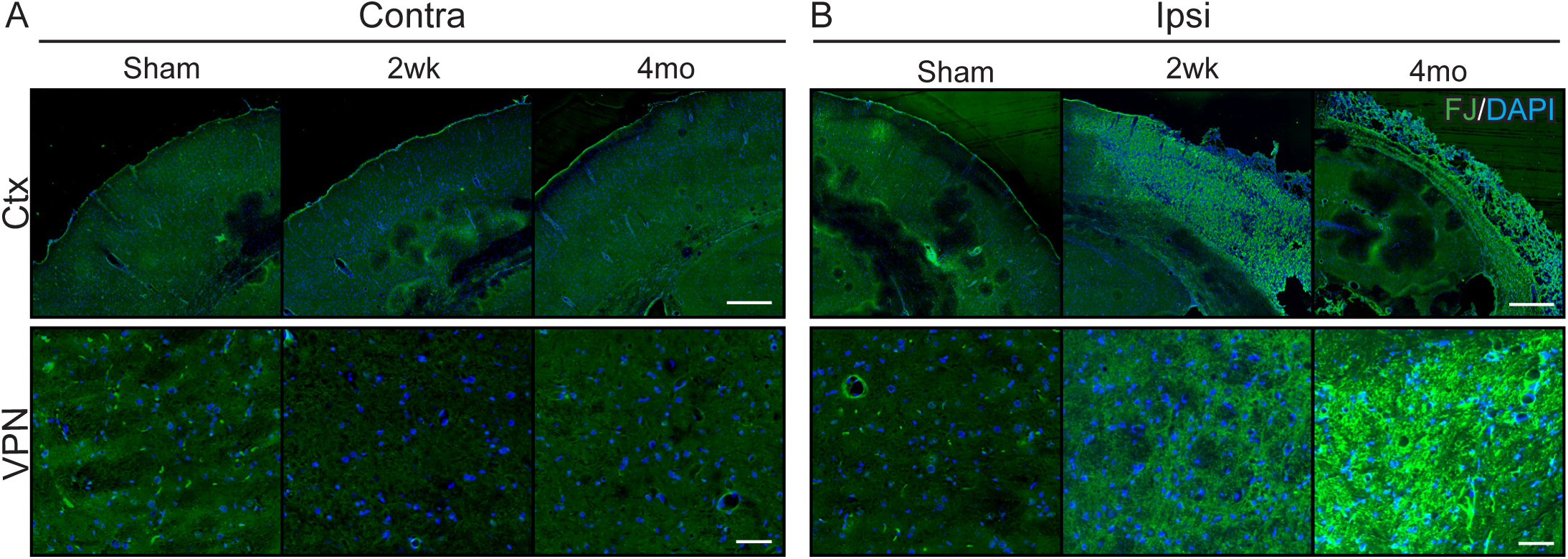
Tissue degeneration in the somatosensory cortex and the ventral posterior nuclei (VPN) of rats at 2 weeks and 4 months after stroke. (**A-B**) Representative image of Fluoro-Jade C (**FJ**) labeling in the (**A**) contralateral and (**B**) ipsilateral cortex. Ctx scale bar, 500 mm and VPN scale bar, 50 mm.

**Supplementary Figure S3.**
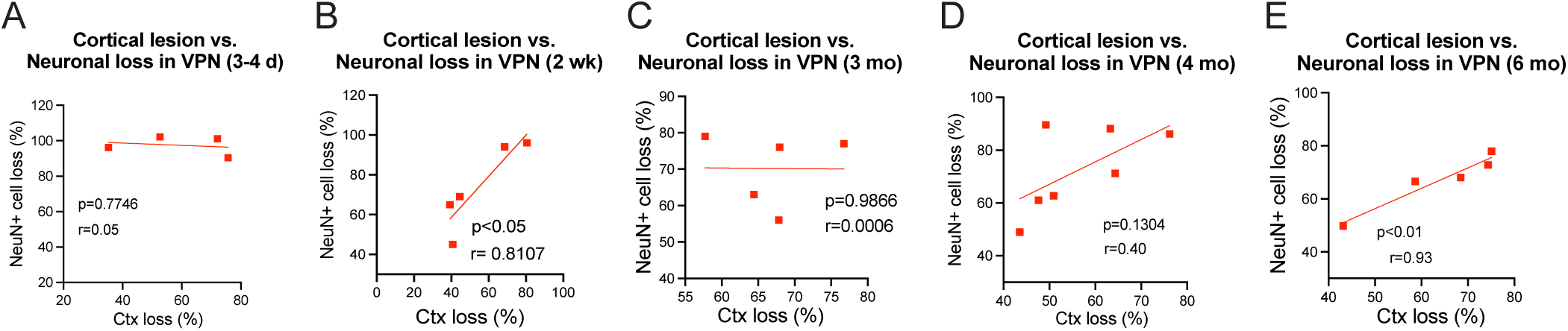
Linear regression analysis showing the correlation between cortical lesion and neuronal loss in the VPN. Correlation was performed at 3-4 days (**A**), 2 weeks (**B**), 3 months (**C**), 4 months (**D)**, and 6 months (**E**) post-stroke. N = 4 - 7 animals per time point. *p<0.05.

**Supplementary Figure S4.**
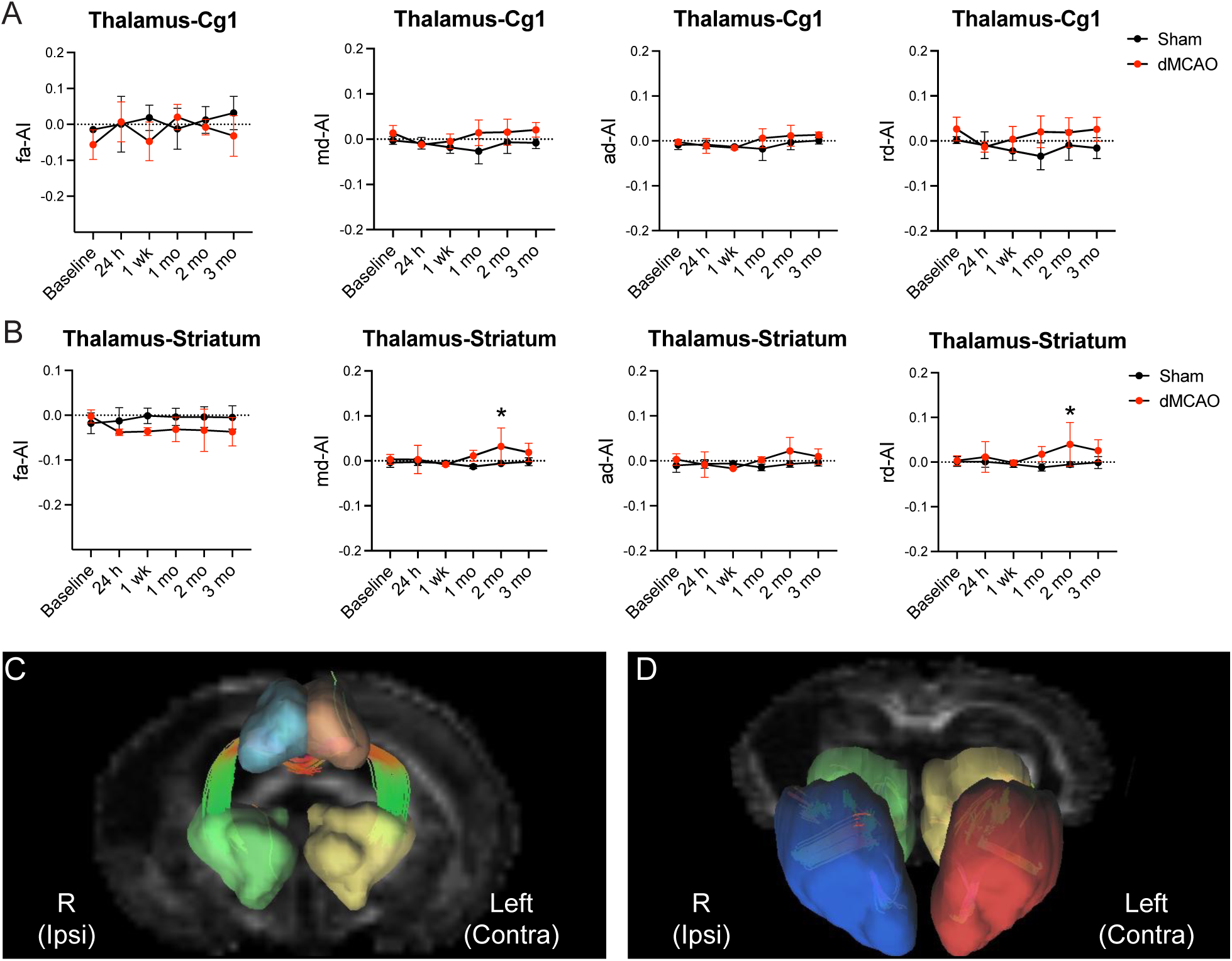
Tractography dynamics of the tracts connecting the ipsilateral thalamus and the stroke-nonaffected regions. (**A-B**) Asymmetry index (AI) of FA, MD, AD, and RD of (**A**) thalamus-primary cingulate cortex (Cg1) and (**B**) thalamus-striatum tracts. (**C-D**) Representative 3D figures demonstrate the fiber tracts connecting the ipsilateral thalamus and (**C**) Cg1 or (**D**) Striatum (Coronal view, with a slight caudal tilt in (D)). Green, right thalamus; Yellow, left thalamus; Sky blue, Cg1; Orange, left Cg1; Blue, right striatum; Red, left striatum. Asymmetry index = (ipsilateral – contralateral)/ (ipsilateral + contralateral). N=4 animals per time point. *p<0.05. Data are presented as mean ± SD.

